# Vegetation structural complexity uniquely captures competition between vascular plants and bryophytes over succession

**DOI:** 10.1101/2024.06.22.600235

**Authors:** Maximilian Hanusch

## Abstract

Vascular plants and bryophytes are among the first colonizers of deglaciated landscapes, thus playing fundamental roles in ecological succession. Understanding biotic interactions between these groups is essential for comprehending the emergence of diverse and complex ecosystems.

Set within a 170-year primary succession gradient in a glacier forefield in the Austrian Alps, this study investigates interactions between vascular plants and bryophytes during early ecosystem development. Data on vegetation structural complexity, obtained through high-resolution 3D-laser scanning, were combined with estimates of local substrate diversity, and the taxonomic and functional composition of vascular plants. This integrated approach allowed us to obtain a mechanistic understanding on the competitive interactions shaping bryophyte diversity over the course of primary succession.

Competitive interactions between vascular plants and bryophytes were uniquely detected through a metric of vegetation structural complexity, while traditional community metrics of vascular plant cover, richness, composition, and functional diversity, failed to reveal these patterns. Path analysis shows that increasing vegetation structural complexity homogenizes local habitat conditions and constrains the realized ecological niche of bryophyte species, while substrate diversity acts as a buffer against vascular plant dominance. This mechanism was quantified using ecological dispersion, a novel concept introduced in this study that estimates the variability of growth optima within a community.

Synthesis: This study highlights the additional value of quantifying the structural properties of plant communities through 3D-laser scanning in explaining ecosystem dynamics. By demonstrating its applicability in grassland ecosystems under field conditions, it underscores the potential of high-resolution remote sensing to advance vegetation structural research. The findings establish vegetation structural complexity as a key factor shaping plant-plant interactions in grasslands, influencing both ecosystem development and biodiversity patterns. These insights may provide valuable guidance for succession-based restoration strategies.

## Introduction

Vegetation structure is fundamental to ecosystem functioning and biodiversity (Loke & Chisholm, 2022). While its importance was recognized early in ecological research (MacArthur et al., 1962; Odum, 1969), recent advances in remote sensing technology have significantly expanded the scope and precision of research in this field (LaRue, Fahey, et al., 2023). Studying the structural properties of plant communities traditionally required destructive field-sampling, which could significantly disturb or damage plant communities (Catchpole & Wheeler, 1992; Zehm et al., 2003). Now, portable 3D-laser scanners enable the precise, non-invasive estimation of vegetation structures across various scales and resolutions, thereby reducing the need for labor-intensive, species-specific or individual-based measurements (Fei et al., 2023; Gámez & Harris, 2022; Zieschank & Junker, 2023). The output of 3D-laser scanning allows for the calculation of various community-level metrics, such as indices of vegetation structural complexity (VSC; Coverdale & Davies, 2023). Similar to how community-weighted means summarize functional traits across an entire community, VSC-metrics represent the three-dimensional distribution of biotic components within an ecosystem. By integrating high-resolution estimates of the physical arrangement of plant structures, VSC metrics provide new opportunities to investigate ecological processes at fine spatial scales (He, Hanusch, Saueressig, et al., 2023; Zieschank & Junker, 2023). Accordingly, indices of VSC may be as valuable and complementary to widely used biodiversity measures, such as taxonomic or functional diversity estimates (Hakkenberg & Goetz, 2021; LaRue, Fahey, et al., 2023; Zhai et al., 2022).

Metrics of VSC represent the occupancy of biotic components within the three-dimensional spatial niche (LaRue, Fahey, et al., 2023), which is closely associated to ecosystem functioning and biotic interactions among organisms (Srivastava, 2006). By expanding three-dimensional habitat space available in an ecosystem, high levels of VSC have been related to increased niche diversity and resource partitioning, making it widely regarded as beneficial for biodiversity (Stein et al., 2014). Accordingly, positive associations between VSC and species diversity have been demonstrated for plant (Coverdale & Davies, 2023), animal (Davies & Asner, 2014; Tews et al., 2004; Traylor et al., 2022), and belowground diversity (Lang et al., 2023). However, most of these insights are derived from forest ecosystems, where trees create strong vertical layering (Davies & Asner, 2014; Gough et al., 2019). Grasslands, in contrast, exhibit limited vertical stratification, as most plants grow within the same vertical layer where they compete for light and other resources. In such environments, herbaceous vascular plants, due to their larger size and greater structural robustness, are expected to have a competitive advantage over smaller-growing plants like bryophytes (Ingerpuu et al., 2005). Thus, the condensed vertical structure of grasslands challenges the assumption that VSC may universally promote the diversity of other organismal groups within a shared habitat.

In structurally complex grasslands, vascular plants develop dense growth forms that create more homogeneous microenvironmental conditions, altering temperature, light availability, and moisture levels, which in turn influence bryophyte communities (Bergamini et al., 2001; Bergamini & Pauli, 2001; Fergus et al., 2017; Guimarães-Steinicke et al., 2021; van Tooren et al., 1990; Werger et al., 2002). Consequently, only bryophyte species with realized ecological niches that align with prevailing habitat conditions are expected to establish and persist (Ridding et al., 2024). Ecological Indicator Values (EIVs) serve as a tool for assessing the realized niche of individual species, representing their growth optima along environmental gradients (Diekmann, 2003). Assessing the dispersion of growth optima across multiple species at the community level, a concept introduced in this study as *ecological dispersion*, provides a measure that estimates the variability of realized niches within a community and is thus indicative of the surrounding habitat conditions (Litvak & Hansell, 1990).

In this study, I test the assumption that vegetation structure is a main driver of vascular plant-bryophyte interactions in grasslands by using 3D-laser scanning along a 170-year primary succession gradient in the Ödenwinkel glacier forefield, Austrian Alps. Successional gradients provide a valuable opportunity to investigate these interactions, as the progressive development of plant communities leads to a gradual increase in VSC over time (Breen & Lévesque, 2006; Cutler et al., 2008; Gavini et al., 2019). Specifically, the following assumptions and hypotheses are tested: 1) VSC captures the distribution of plant components in three dimensions, thereby providing ecological information that extends beyond traditional two-dimensional or individual-based vegetation metrics. VSC is thus expected to contain additional, ecologically meaningful information that is not fully captured by traditional two-dimensional metrics of vascular plant communities, namely species composition, plant cover, as well as taxonomic and functional diversity. 2) Over grassland succession, increasing VSC is expected to constrain bryophyte diversity by reducing available niche space through competitive exclusion and habitat homogenization.

## Methods

### Study design

The study was conducted in the long-term ecological research platform Ödenwinkel which was established in 2019 in the Hohe Tauern National Park, Austria (Dynamic Ecological Information Management System – site and dataset registry: https://deims.org/activity/fefd07db-2f16-46eb-8883-f10fbc9d13a3, last access: May 2025; Junker *et al*., 2020). A total of *n* = 135 permanent plots were established within the glacier forefield of the Ödenwinkelkees, which was covered by ice at the latest glacial maximum in the Little Ice Age around 1850. The plots represent a successional gradient spanning over 1.7km in length at an altitude ranging from 2070 to 2170 m a.s.l. with a minimum distance of 5 m between plots. Plots were defined as squares with an area of 1x1 m and were all oriented in the same cardinal direction. Detailed descriptions of the research platform, exact plot positions and details on the surrounding environment can be found in Junker et al. (2020) and Hanusch et. al (2022). Fieldwork was conducted with permission from the governing authority of Land Salzburg (permit no. 20507-96/45/7-2019).

### Biotic and abiotic sampling

Plot age was estimated according to the plots’ relative position compared with historical records of eight deglaciation periods (year 1850, 1890, 1929, 1969, 1977, 1985, 1998, 2013; Junker et al., 2020). To assess the substrate composition of the plots, the percentage cover of bare ground, litter, scree and solid rock were estimated.

During the 2019 field season, all bryophyte species on each plot were identified and recorded (*n* = 42 in 135 plots), noting their coverage at a 0.1% resolution. This was accomplished by using 1m² vegetation relevés subdivided into 100 squares, each measuring 10 × 10 cm. Likewise, all vascular plant species (n = 93 across 135 plots) were recorded. Both communities were sampled in the same plots, which allows a direct comparison of potential interactions between both groups (He, Hanusch, Ruiz-Hernández, et al., 2023; R. R. Junker et al., 2021). Furthermore, based on estimates of leaf area and weight, specific leaf area, and growth height vascular plant functional traits were calculated as described in Junker *et al*. (2020) and He, Hanusch, Saueressig, *et al*. (2023). A full dataset, i.e. no NA value for any variable, was used for this study representing a subset of *n* = 107 out the total *n* = 135 plots.

### 3D-laser scanning and vegetation structural complexity

In the vegetation period 2021 (from 16 July to 23 July), an on-site plot scanning in the field was conducted with a multispectral 3D-laser-scanner ‘PlantEye’ (Phenospex, F500, Heerlen, the Netherlands; Supplementary Figure 1). The plant eye is equipped with an active sensor that projects a near-infrared (940 nm) laser line vertically onto the plant canopy (Kjaer & Ottosen, 2015; Maphosa et al., 2017). Each scan has a resolution of < 1 pixel*mm ^–1^ and the data points of each scan are merged to generate a 3D point cloud. From these clouds, a set of community level structural properties can be estimated that have been shown to correlate well with real world manual measurements (Zieschank & Junker, 2023). As colonization and growth rate of plant species tend to remain relatively stable over short timescales, even in dynamic environments like glacier forefields (Bayle et al., 2023), considerable shifts in vascular plant presence, abundance, or structural characteristics are not expected to have occurred between the timepoints of vegetation sampling and scanning. Settings of software used to calculate community features followed He *et al*. (2023), for a detailed description of the set-up and data obtained, see Zieschank & Junker (2023). In this study digital community measurements of biomass (BM), leaf area index (LAI) and growth height-corrected light penetration depth (LPD) were incorporated: LPD is a measure for the depth to which light can penetrate through the vegetation layer. The plant eye calculates this value as the distance between the on average highest 10 % and on average lowest 20 % of the vegetation layer that the light can penetrate through. As a result, the uncorrected, raw measurements of LPD are not independent of the vegetation’s growth height. Thus, a growth height-corrected LPD-index was calculated by subtracting LPD from the on average highest 10 % of the vegetation layer. This index is an estimate of the height of the lowest and most dense 20 % of the vegetation. Low growth height-corrected LPD values indicate a shorter, scattered vegetation; high values indicate a higher, more dense vegetation. The LAI measures the density of vascular plant leaves in a given area, representing the ratio of one-sided leaf area to ground area (Coverdale & Davies, 2023); i.e. high values indicate a denser vegetation with a high number of leaves, low numbers indicate a sparse vegetation with a low number of leaves. The estimates of BM represent a digital approximation of the standing biomass, and is calculated as the product of height and 3D leaf area.

VSC is not directly observable and there is still no consensus over the definition of structural complexity or how to measure it (Atkins et al., 2018; Loke & Chisholm, 2022). In this study VSC is quantified as a univariate variable generated through a principal component analysis (PCA; R package ’vegan’ v2.6-4; Oksanen et al., 2019) conducted on normalized estimates of BM, LAI, and growth height-corrected LPD derived from 3D-laser-scanning. The first principal component (PC1), which explained 84.0% of the variance, was used as the structural complexity metric.

### Traditional metrics of vascular plant communities

To examine whether VSC adds additional information or is redundant to traditional vascular plant community metrics in explaining bryophyte diversity, four additional metrics were assessed: vascular plant species cover, species richness, taxonomic composition, and functional diversity. Species richness represents the total number of vascular plant species per plot, taxonomic composition was summarized using the first principal component (15% explained variance) from a PCA of vascular plant species composition. Functional diversity was quantified as functional divergence using the fd_fdiv command in the fundiversity-package (v1.1.1, R) based on trait measurements of leaf area and weight, specific leaf area, and growth height. High values of functional divergence suggest functional niche differentiation, potentially reflecting similar dimensions of spatial niche occupancy as those captured by measures of VSC. Species cover was measured as the percentage area covered by vascular plants on each plot. Pearson correlations were conducted to evaluate the relationships among these metrics and structural complexity, as well as with bryophyte diversity.

### Calculation of ecological dispersion

Ecological dispersion provides a novel framework to examine shifts in a community’s realized ecological niche driven by changing habitat conditions. Ecological dispersion, in contrast to the traditional use of EIVs to track shifts in single environmental factors, captures the variability of species’ preferences across multiple factors, providing an estimate of habitat heterogeneity. To calculate ecological dispersion of bryophyte communities, EIV data for bryophytes were obtained from the 2017 updated version of BryoATT (Hill et al., 2007). In total, EIV assignments for L (light), R (pH), F (humidity), and N (nutrients) were successfully made for *n* = 32 bryophyte species; i.e. 76% of all recorded bryophyte taxa.

To evaluate whether the selected EIVs represent ecologically meaningful variables, I assessed the relationship of the abundance-weighted mean EIV of all bryophyte species occurring in a plot with VSC using Spearman’s rank correlation. Spearman correlation was chosen because a linear relationship was not expected, as community-weighted means can be strongly influenced by the presence of species with distinct EIV values, potentially causing abrupt shifts in community means. Community-weighted means of F and N showed a significant positive relationship with VSC (*rho_F_* = 0.19, *p_F_* = 0.05; *rho_N_* = 0.23, *p_N_* = 0.01), while community-weighted means of L showed a significant negative relationship (*rho_L_* = -0.23, *p_L_* = 0.01), and no significant relationship with R (*rho_R_* = -0.13, *p_R_* = 0.17).

In a next step, ecological dispersion was calculated as follows: First the ordinal EIV for each species were transformed into a continuous form by building a multidimensional functional space using principal coordinate analysis (ape v5.7-1 package, R) based on Euclidean-distances of the four EIV-values (De Bello et al., 2021; Maire et al., 2015). Second, ecological dispersion was calculated based on the axes coordinate of each species in each of the four dimensions by using fd_fdis command in the fundiversity-package (v1.1.1, R). High ecological dispersion indicates a heterogeneous habitat that supports species with diverse ecological requirements, whereas low dispersion reflects homogeneous conditions favoring species with similar ecological requirements.

### Path analysis

Estimating biotic interactions from observational field data remains inherently challenging, particularly in the context of ecological succession (Blanchet et al., 2020; B. J. W. Chen et al., 2022; Hanusch et al., 2023, 2024). To test the hypothesis that VSC drives competition between vascular plants and bryophytes, a path analysis was conducted using the R package lavaan (v0.6-1.7; Rosseel, 2012; Tsukahara, 2023). Path analysis was selected for its ability to disentangle direct and indirect effects among multiple interacting variables in predeveloped theoretical models and obtain estimates of the explained variance (R²) by predictors. The model included key variables hypothesized to influence bryophyte diversity: Time since deglaciation represents the increasing likelihood of heterogeneous dispersal events over time (Ficetola et al., 2024; Hanusch et al., 2022a). VSC served as a proxy for competitive interactions, capturing how the physical structure of vascular plant communities may restrict bryophytes growing within the plant layer. Additionally, substrate diversity, calculated as the Shannon diversity of relative cover of substrate types (bare ground, litter, scree, and solid rock), was included as a univariate variable to represent environmental heterogeneity. Substrate diversity showed a significant correlation with bryophyte composition (Mantel-test: Mantel r = 0.26, p < 0.001), highlighting its relevance to this plant group. As the study was conducted along a linear successional gradient in a glacier forefield, spatial distance was highly correlated with time since deglaciation (Mantel-test: Mantel r = 0.88, p < 0.001) and was therefore excluded from the model to avoid multicollinearity. The path analysis aimed to determine whether changes in bryophyte diversity are primarily shaped by environmental heterogeneity, dispersal processes, or interactions with vascular plants. All variables were standardized by z-transformation by subtracting the mean and dividing through the standard deviation.

In the model, time since deglaciation is included as an exogenous variable, exerting direct effects on VSC, substrate diversity, and bryophyte ecological dispersion. VSC and substrate diversity, in turn, have direct impacts on bryophyte ecological dispersion. Additionally, the model examines indirect effects of time since deglaciation, VSC, and substrate diversity on bryophyte diversity, mediated through ecological dispersion.

To evaluate whether VSC provides additional information compared to traditional plant community metrics, four alternative model structures were developed. In each model VSC was replaced with one of the traditional plant community metrics: vascular plant cover, species richness, species composition (first axis from PCA), or functional divergence.

## Results

### Comparison between vascular plant community metrics and their relationship with bryophyte diversity

The VSC-metric showed a significant negative relationship with bryophyte diversity (*t* = -2.00, *df* = 104, *r^2^*= 0.04, *p* = 0.04), which was not observed with other vascular plant community metrics (Figure 1). Plant cover exhibited a non-linear relationship with bryophyte diversity (*t* = -2.68, *df* = 103, *r^2^* = 0.07, *p* < 0.01), while plant species richness showed a marginally significant positive linear relationship (*t* = 1.80, *df* = 104, *r^2^* = 0.03, *p* = 0.06). Neither species composition (*p* = 0.64) nor functional divergence (*p* = 0.50) demonstrated a significant linear relationship with bryophyte diversity. Among the vascular plant community metrics, VSC showed the highest correlation with plant cover (*r* = 0.72, *p* < 0.01, Figure 1F), followed by species composition (*r* = 0.58, *p* < 0.01), species richness (*r* = 0.38, *p* < 0.01), and the lowest correlation with functional divergence (*r* = 0.23, *p* = 0.02).

**Figure 1:**
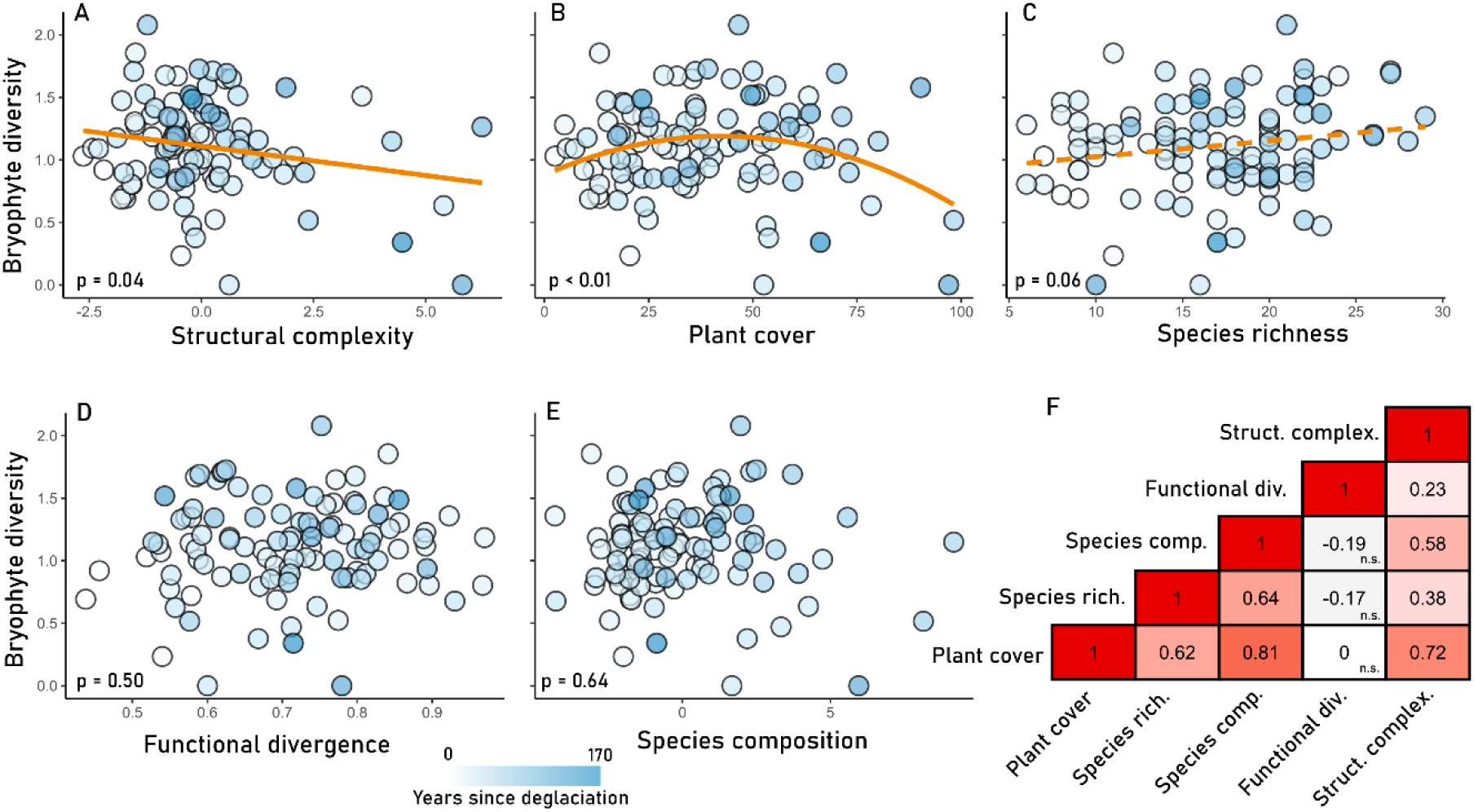
Relationship between bryophyte diversity (BD) and five metrics of the vascular plant community growing in the same habitat across *n* = 107 plots along a primary succession gradient spanning 170 years. Darker points indicate older plots. (A) significant negative linear relationship between vascular plant structural complexity, estimated as the first axis of a PCA on digital biomass, leaf area index, and height-corrected light penetration depth and BD. (B) significant non-linear relationship (quadratic fit) between vascular plant cover and BD. (C) marginally significant positive linear relationship between vascular plant richness and BD. (D) non-significant relationship between vascular plant functional diversity and BD. (E) non-significant relationship between vascular plant community composition and BD. (F) Pearson correlation heatmap showing relationships among the four plant community metrics and VSC.

### Path analysis

The path analysis provided good model fit (Model fit: p_chi-square_ > 0.05; CFI > 0.95; TLI > 0.9; SRMR < 0.05; RMSEA = 0.08 with lower bound of confidence interval = 0) and revealed significant effects of all exogenous variables on bryophyte ecological dispersion (Figure 2). Time since deglaciation (*β* = 0.25, *p* = 0.01) and substrate diversity (*β* = 0.21, *p* = 0.03) positively affected bryophyte ecological dispersion. Conversely, VSC showed a significant negative correlation with ecological dispersion (*β* = -0.29, *p* < 0.01). Time since deglaciation had a negative effect on substrate diversity (*β* = -0.26, *p* < 0.01) but a positive effect on VSC (*β* = 0.42, *p* < 0.001). Bryophyte diversity was positively affected by bryophyte ecological dispersion (*β* = 0.83, *p* < 0.001) and experienced positive indirect effects by substrate diversity (*β* = 0.17, *p* = 0.03) while VSC had a negative indirect effect (*β* = -0.24, *p* < 0.01). The total indirect effects of time since deglaciation on ecological dispersion (i.e. temporal processes including effects mediated through substrate diversity and VSC) were non-significant (*β* = 0.07, *p* = 0.40) indicating that several simultaneous processes may act over time that offset each other. Thus, the indirect effects of time since deglaciation on bryophyte diversity were separately calculated for the effects mediated through substrate diversity (*β* = -0.04, *p* = 0.09), increasing VSC over time (*β* =-0.10, *p* = 0.02), and time since deglaciation in isolation (i.e. temporal processes without effects of substrate diversity and VSC; *β* = 0.21, *p* = 0.02). The four alternative models, in which VSC was replaced with one of the four other vascular plant community metrics (vascular plant species richness, functional diversity, cover, and composition) showed no significant effects of these variables on bryophyte diversity or associated ecological dispersion, which highlights VSC as the primary factor influencing bryophyte diversity (Supplementary tables 1-5). These relationships are synthesized in the conceptual framework shown in Figure 3, which illustrates how biotic and abiotic drivers interact to shape bryophyte diversity through their effects on ecological dispersion.

**Figure 2:**
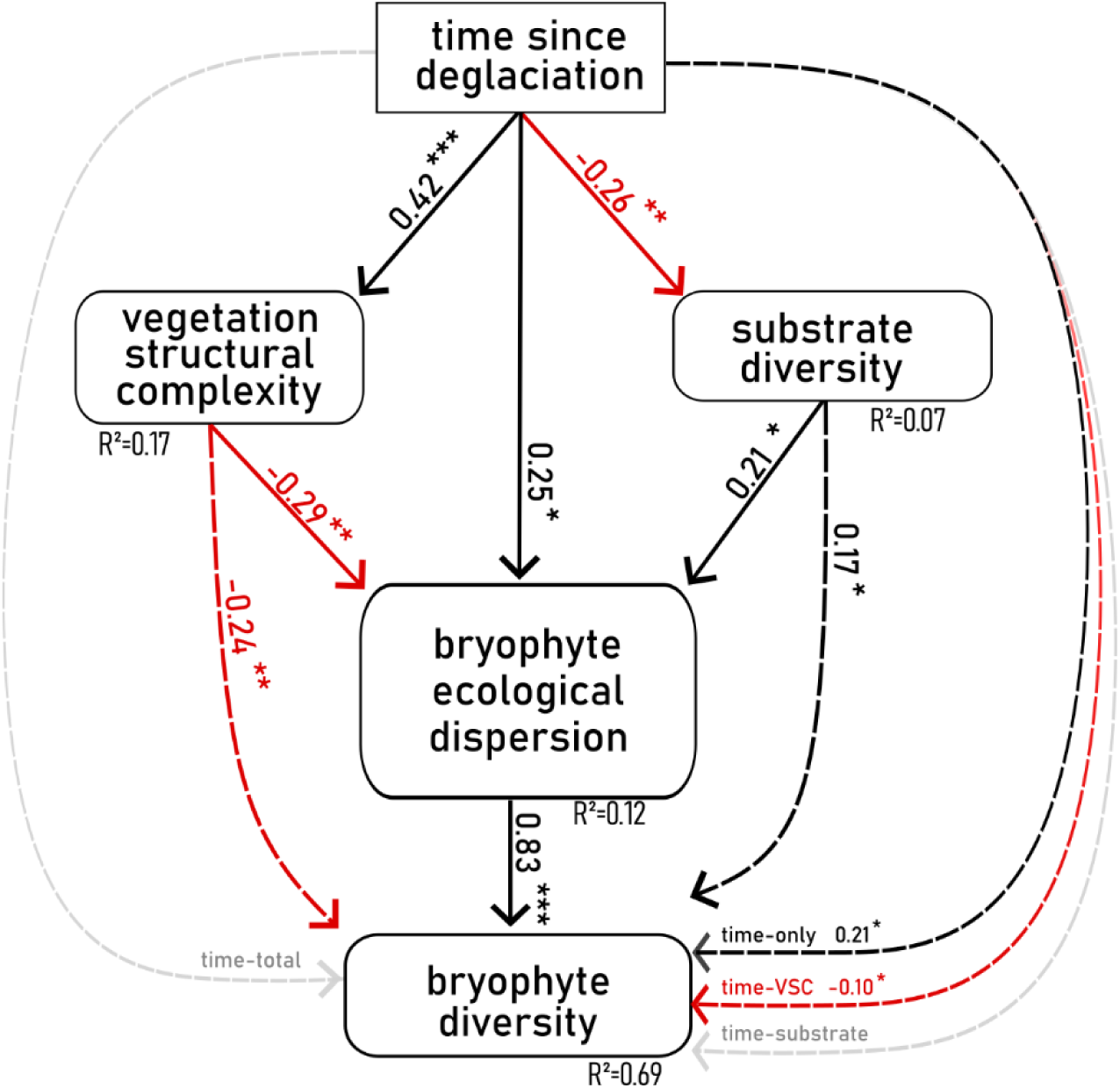
Path analysis estimating the effects of time since deglaciation, substrate diversity and vegetation structural complexity (VSC) on bryophyte ecological dispersion and bryophyte diversity. The path diagram presents direct effects as solid lines. Indirect effects are represented as dotted lines and were calculated through mediation analysis. Time-total refers to the total indirect effects of time since deglaciation including the mediated effects through both VSC and substrate diversity. Time-only refers to the indirect effects of time excluding mediation through VSC and substrate diversity. Time-VSC and Time-substrate refer to the indirect effects mediated solely through VSC and substrate diversity, respectively. Standardized path coefficients are shown for significant relationships, with red coloration indicating a negative and black coloration indicating a positive effect. Asterisks indicate significance levels (* = p ≤ 0.05; ** = p ≤ 0.01; *** = p ≤ 0.001).

**Figure 3:**
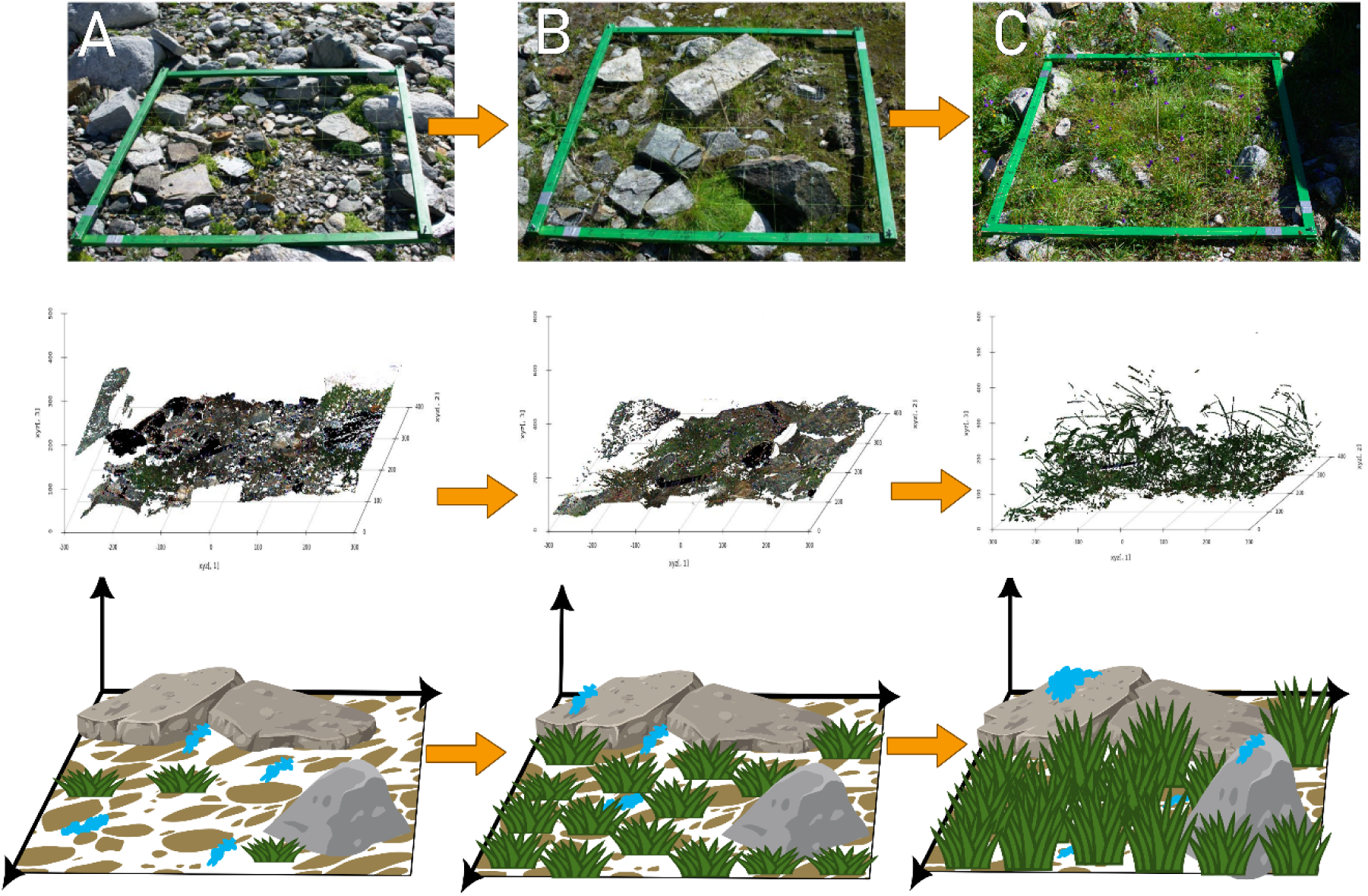
Conceptual synthesis of the mechanisms shaping bryophyte community assembly over grassland succession as inferred from model results. Each triplet, comprising a photograph, a 3D laser scan of vegetation, and a conceptual 3D sketch, represents a plot with increasing successional age and vascular plant structural complexity (VSC). Bryophyte patches are highlighted in blue in the conceptual sketches. Over succession, increasing VSC leads to denser plant structures that reduce and homogenize the available habitat for bryophytes. However, substrate weathering and heterogeneous microhabitats such as exposed rocks provide refuges for bryophyte persistence. These effects are particularly evident in plots with high substrate diversity. (A) A plot with low VSC, characterized by a high proportion of open substrate and exposed rock surfaces (∼6 years since deglaciation). (B) A plot with moderate VSC, where vascular plant density has increased (∼35 years since deglaciation). (C) A plot with high VSC, where vascular plants dominate and bryophytes persist mainly in isolated microhabitats (∼100 years since deglaciation). Sketches published by OpenClipArt, permission granted under CC0 1.0 Universal.

## Discussion

The results indicate that competition with vascular plants plays an important role in shaping bryophyte diversity over grassland succession following glacial retreat. Increasing vegetation structural complexity appears to homogenize local habitat conditions, thereby narrowing the ecological niches available to bryophytes. Traditional metrics such as vascular plant cover, species composition, richness, and functional divergence did not capture these interactions. These results highlight vegetation structural complexity as a key factor influencing biodiversity patterns.

### Vascular plants shape bryophyte succession through biotic interactions

The path analysis revealed that VSC negatively affects both bryophyte diversity and ecological dispersion along the successional gradient, resulting in less diverse bryophyte communities with more homogeneous ecological preferences in structurally complex habitats. These competitive interactions are likely driven by increasingly dense and complex plant structures that alter microclimatic conditions, as well as intensified competition for light and space (Alatalo et al., 2020; Bergamini et al., 2001; Schwager et al., 2025). Although no direct measurements of light availability were conducted on-site, light-penetration depth, one of the variables used in constructing the VSC metric, serves as an indirect proxy for light availability. Additionally, the observed negative correlation between VSC and bryophyte CWM-estimates of light-EIV, alongside a positive correlation with moisture-EIV, suggests that vascular plants outcompete bryophytes by modifying the habitat towards more moist and shaded conditions. This change in habitat conditions likely restricts the available niche space, favoring bryophyte species adapted to low-light, high-moisture environments.

Despite ongoing competition, bryophyte diversity remained higher on plots with more diverse substrate compositions and on older plots. Bryophytes in glacier forefields are generally not considered dispersal-limited (Cichini et al., 2011), suggesting that the positive effect of time since deglaciation cannot be solely attributed to slow dispersal processes, which are more typical of vascular plant communities (Cantera et al., 2024; Hanusch et al., 2022a; Ulrich et al., 2016). Instead, site-specific processes, such as substrate weathering, likely play a role by facilitating opportunistic and rapid colonization of newly available niches (Prach & Walker, 2020). The positive effect of substrate diversity on bryophyte diversity aligns with previous studies, confirming that the available substrate is a key driver of bryophyte community composition and diversity during succession (e.g. Hylander *et al*., 2005; Caruso & Rudolphi, 2009; Chen *et al*., 2017; Spitale, 2017). Accordingly, bryophyte succession in alpine grasslands is driven by a combination of interrelated processes. The successful colonization of microhabitats within a homogenous environment helps buffer bryophyte diversity against competition with vascular plants over time. This highlights the importance of habitat heterogeneity in maintaining species coexistence and underscores the need to consider microhabitat dynamics when studying plant community assembly and resilience in changing ecosystems.

### Ecological dispersion as a measure for environmental changes

In this study, EIVs were used to estimate the variability in growth optima of bryophyte communities by calculating their ecological dispersion (Laliberté et al., 2014; Litvak & Hansell, 1990). While EIV values are typically averaged across communities to describe growth optima for individual environmental factors, ecological dispersion captures the breadth and heterogeneity of growth optima across multiple ecological dimensions. This approach provides a more nuanced understanding of local niche diversity and habitat heterogeneity, and allows insights into how niche differentiation may structure species coexistence within communities. It is well recognized that both biotic and abiotic factors influence the variability of functional strategies at the community level (Bruelheide et al., 2018; Díaz et al., 1998; Neyret et al., 2024). Similarly, in this study, substrate diversity and VSC are important biotic and abiotic factors that shape local habitat conditions and favor bryophyte species with growth optima suited to these environments, thus driving changes in the community’s ecological dispersion (Sterck et al., 2011). Despite certain limitations of EIVs, such as their ordinal nature and sensitivity to spatial or temporal scale (Berg et al., 2017; Diekmann, 2003; Ewald, 2003, 2009), ecological dispersion proved to be a valuable proxy in identifying these filtering mechanisms. Thus, assessments of ecological dispersion could serve as a powerful complement to existing approaches that have applied EIV values of bryophytes for detecting environmental changes (Delgado & Ederra, 2013; Mayo de la Iglesia et al., 2024; Pakeman et al., 2019).

### Adopting a “three-dimensional perspective” for ecological research

A key advantage of laser scanning techniques is their ability to estimate properties of plant communities in three dimensions, explicitly accounting for the spatial arrangement of physical plant structures across both horizontal and vertical dimensions (LaRue, Fahey, et al., 2023). This capability represents a significant improvement over traditional vascular plant community metrics, which are typically derived from two-dimensional assessments based on plant cover or individual-based estimates of abundance. Such two-dimensional approaches often fail to capture the inherently three-dimensional aspects of plant communities that strongly influence biodiversity patterns within an ecosystem (Gámez & Harris, 2022). In this study, VSC was the only metric that showed a significant negative correlation with bryophyte diversity, due to its capability to account for the physically occupied niche spaces in ecosystems (LaRue, Knott, et al., 2023). The spatial arrangement of plants, varying in size and structure, creates distinct patterns of niche occupation along both horizontal and vertical axes, particularly for resources such as light and space (Forrester et al., 2018). Size-asymmetric competition for these resources is a well-documented driver of grassland biodiversity, as larger plants overshadow smaller ones, restricting their access to essential resources and limiting their growth (Eskelinen et al., 2022; Gruntman et al., 2017; Newman, 1973; Weiner, 1990). This form of competition primarily operates along the vertical axis of the three-dimensional habitat space, as it is fueled by the directional supply of light and thus may fail to be acknowledged by traditional two-dimensional metrics. In this study, VSC integrated key structural properties of plant communities, specifically light penetration depth, leaf area index, and digital biomass. While light penetration depth and leaf area index act as proxies for light availability to understory species, digital biomass serves as a proxy for spatial competition by quantifying the three-dimensional volume occupied by vegetation (Grace, 1999; Zieschank & Junker, 2023). Building on these insights, integrating VSC into ecological research provides a promising approach for understanding the structural drivers of plant-plant interactions and their ecological consequences.

Although traditional vegetation metrics may not capture specific relationships revealed by VSC, they provide complementary rather than redundant information. Correlations between structural complexity and traditional metrics indicate that while some measures are closely related, others reflect distinct aspects of vascular plant communities. Adopting a “three-dimensional perspective” broadens ecosystem analyses beyond traditional two-dimensional assessments of plant communities by acknowledging their volumetric properties; an approach that could provide a more comprehensive understanding of ecosystem processes and dynamics (B. J. W. Chen et al., 2022; Davies & Asner, 2014). As demonstrated in this study, VSC is a strong and unique predictor of bryophyte responses to environmental and biotic changes, capturing dynamics that remain undetected by traditional vegetation metrics. To fully understand the mechanisms shaping biodiversity patterns, ecologists must carefully consider which metrics they use, as each provides distinct yet complementary insights into ecosystem processes. Integrating structural properties into ecological research may not only refine our understanding of plant-plant interactions but may also open new avenues for exploring how habitat complexity drives community assembly and ecosystem resilience in a changing world.

## Conclusion

This research contributes to the ongoing integration of VSC as a key component of biodiversity studies (Coverdale & Davies, 2023; LaRue, Knott, et al., 2023). Building on the hypothesis that VSC captures ecologically meaningful information beyond traditional two-dimensional metrics, this study demonstrates how three-dimensional plant architecture shapes plant–plant interactions and biodiversity patterns. The findings support the expectation that increasing VSC during grassland succession can constrain bryophyte diversity by homogenizing habitat conditions and limiting niche space. Nonetheless, bryophytes persist by colonizing diverse microhabitats that emerge over time. Advanced laser-scanning methods and VSC modeling provide powerful tools for understanding species coexistence and ecosystem dynamics. By integrating structural metrics with conventional vascular plant community assessments, researchers can gain a more holistic view of ecosystem functioning. Such approaches pave the way for more effective conservation, succession-based restoration strategies, and a deeper understanding of ecological complexity.

## Contributions

Maximilian Hanusch designed the study, conducted field work, performed the processing and analysis of the data and wrote the manuscript.

## Data availability

Raw floristic and environmental data is available as a Mendeley Data repository under DOI: 10.17632/xkv89tbftc.1 (Hanusch et al., 2022b). The dataset on multispectral 3D plant scanning be found under DOI: 10.17632/ztmwmbg9pd.3 (R. Junker et al., 2023).

## Acknowledgements

I thank Robert R. Junker and Manuel Pitzer for fruitful discussions on the ecological relevance of vegetation structural complexity and for their support in interpreting the measurements obtained through 3D laser scanning. I am also grateful to the Hohe Tauern National Park Salzburg administration and the Rudolfshütte for their organizational and logistical support, and to the governing authority Land Salzburg for granting the research permit (permit no. 20507-96/45/7-2019). I further thank an anonymous reviewer for valuable comments that helped improve the quality of the manuscript. This research has been supported by the Austrian Science Fund (F.W.F.), which provided funding to Robert R. Junker (grant no. Y1102).

## Competing interests

The author declares no competing interests.

## Supplementary Information

**Supplementary Figure 1:**
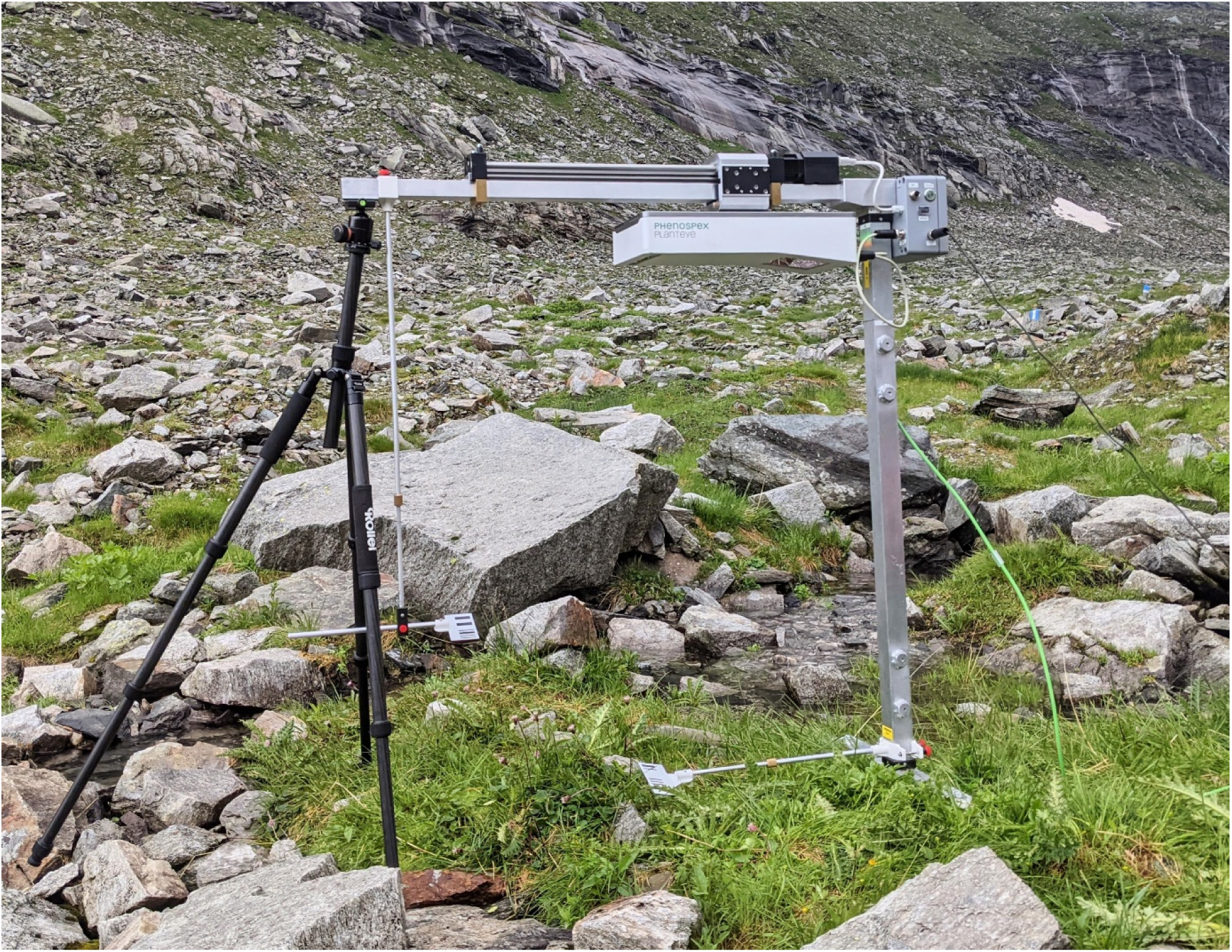
Multispectral 3D-laser-scanning using a customized PlantEye setup (Phenospex, F500, Heerlen, the Netherlands) in the field.

## Notes

### Competing Interest Statement

The authors have declared no competing interest.

### Summary of Updates

Updated results; new analysis and figures;

