## Supplemental tables for "Vegetation structural complexity uniquely captures competition between vascular plants and bryophytes over succession"

Supplementary table 1: Summary of the path analysis of the effects of time since deglaciation, vegetation structural complexity and substrate diversity on bryophyte ecological dispersion and diversity (final model).

| Model Significance | | | |  |  |  |  |
| --- | --- | --- | --- | --- | --- | --- | --- |
| **N** | **χ^2^** | **df** | **p** |  |  |  |  |
| 106 | 6.559 | 4 | 0.161 |  |  |  |  |
| Model Fit | | | | | | |  |
| **CFI** | **RMSEA** | **90% CI** | **TLI** | **SRMR** | **AIC** | **BIC** |  |
| 0.984 | 0.078 | 0.000 — 0.180 | 0.96 | 0.025 | 1064.332 | 1101.62 |  |
| Regression Paths | | | | | | | |
| **Predictor** | **DV** | **Standardized** | | | | | |
|  |  | **β** | **95% CI** | **sig** | **SE** | **z** | **p** |
| (c*f) | age.div.env | −0.053 | −0.113 — 0.007 |  | 0.031 | −1.733 | 0.083 |
| (a*d) | age.div.plant | −0.119 | −0.212 — −0.027 | * | 0.047 | −2.529 | 0.011 |
| (c*f)+(a*d)+(e*g) | age.div.total | 0.039 | −0.124 — 0.202 |  | 0.083 | 0.467 | 0.640 |
| (c*f*g) | age.obs.env | −0.044 | −0.094 — 0.006 |  | 0.026 | −1.725 | 0.085 |
| (e*g) | age.obs.isolated | 0.211 | 0.047 — 0.375 | * | 0.084 | 2.526 | 0.012 |
| (a*d*g) | age.obs.plant | −0.099 | −0.177 — −0.022 | * | 0.04 | −2.503 | 0.012 |
| (a*d*g)+(c*f*g)+(e*g) | age.obs.total | 0.068 | −0.089 — 0.225 |  | 0.08 | 0.849 | 0.396 |
| age | env | −0.258 | −0.433 — −0.083 | ** | 0.089 | −2.893 | 0.004 |
| (f*g) | env.ind | 0.171 | 0.018 — 0.324 | * | 0.078 | 2.196 | 0.028 |
| obs.mos | mos.div | 0.831 | 0.772 — 0.890 | *** | 0.03 | 27.638 | 0.000 |
| age | obs.mos | 0.254 | 0.059 — 0.450 | * | 0.1 | 2.555 | 0.011 |
| env | obs.mos | 0.206 | 0.024 — 0.388 | * | 0.093 | 2.216 | 0.027 |
| pc.struc | obs.mos | −0.287 | −0.478 — −0.097 | ** | 0.097 | −2.956 | 0.003 |
| age | pc.struc | 0.416 | 0.265 — 0.566 | *** | 0.077 | 5.414 | <0.001 |
| (d*g) | plant.ind | −0.239 | −0.400 — −0.078 | ** | 0.082 | −2.909 | 0.004 |
| * p < .05; ** p < .01; *** p < .001 | | | | | | | |

Supplementary table 2: Summary of the path analysis of the effects of time since deglaciation, vascular plant species richness and substrate diversity on bryophyte ecological dispersion and diversity (model not used).

| Model Significance | | | |  |  |  |  |
| --- | --- | --- | --- | --- | --- | --- | --- |
| **N** | **χ^2^** | **df** | **p** |  |  |  |  |
| 106 | 2.888 | 3 | 0.409 |  |  |  |  |
| Model Fit | | | | | | |  |
| **CFI** | **RMSEA** | **90% CI** | **TLI** | **SRMR** | **AIC** | **BIC** |  |
| 1 | 0 | 0.000 — 0.161 | 1.002 | 0.029 | 1061.694 | 1101.646 |  |
| Regression Paths | | | | | | | |
| **Predictor** | **DV** | **Standardized** | | | | | |
|  |  | **β** | **95% CI** | **sig** | **SE** | **z** | **p** |
| (c*f) | age.div.env | −0.058 | −0.121 — 0.005 |  | 0.032 | −1.803 | 0.071 |
| (a*d) | age.div.plant | 0.064 | −0.037 — 0.166 |  | 0.052 | 1.240 | 0.215 |
| (c*f)+(a*d)+(e*g) | age.div.total | 0.095 | −0.069 — 0.260 |  | 0.084 | 1.134 | 0.257 |
| (c*f*g) | age.obs.env | −0.048 | −0.105 — 0.009 |  | 0.029 | −1.662 | 0.096 |
| (e*g) | age.obs.isolated | 0.089 | −0.094 — 0.272 |  | 0.093 | 0.951 | 0.342 |
| (a*d*g) | age.obs.plant | 0.054 | −0.035 — 0.143 |  | 0.045 | 1.191 | 0.234 |
| (a*d*g)+(c*f*g)+(e*g) | age.obs.total | 0.094 | −0.068 — 0.257 |  | 0.083 | 1.137 | 0.255 |
| age | env | −0.258 | −0.433 — −0.083 | ** | 0.089 | −2.893 | 0.004 |
| (f*g) | env.ind | 0.187 | 0.010 — 0.365 | * | 0.09 | 2.072 | 0.038 |
| age | mos.div | 0.106 | −0.107 — 0.319 |  | 0.109 | 0.975 | 0.329 |
| env | mos.div | 0.224 | 0.039 — 0.410 | * | 0.095 | 2.371 | 0.018 |
| plant.ric | mos.div | 0.134 | −0.073 — 0.342 |  | 0.106 | 1.267 | 0.205 |
| mos.div | obs.mos | 0.836 | 0.454 — 1.219 | *** | 0.195 | 4.283 | <0.001 |
| (d*g) | plant.ind | 0.112 | −0.069 — 0.293 |  | 0.092 | 1.214 | 0.225 |
| age | plant.ric | 0.481 | 0.343 — 0.618 | *** | 0.07 | 6.846 | <0.001 |
| * p < .05; ** p < .01; *** p < .001 | | | | | | | |

Supplementary table 3: Summary of the path analysis of the effects of time since deglaciation, vascular plant composition and substrate diversity on bryophyte ecological dispersion and diversity (model not used).

| Model Significance | | | |  |  |  |  |
| --- | --- | --- | --- | --- | --- | --- | --- |
| **N** | **χ^2^** | **df** | **p** |  |  |  |  |
| 106 | 0.628 | 3 | 0.89 |  |  |  |  |
| Model Fit | | | | | | |  |
| **CFI** | **RMSEA** | **90% CI** | **TLI** | **SRMR** | **AIC** | **BIC** |  |
| 1 | 0 | 0.000 — 0.075 | 1.052 | 0.013 | 1066.381 | 1106.333 |  |
| Regression Paths | | | | | | | |
| **Predictor** | **DV** | **Standardized** | | | | | |
|  |  | **β** | **95% CI** | **sig** | **SE** | **z** | **p** |
| (c*f) | age.div.env | −0.051 | −0.109 — 0.007 |  | 0.03 | −1.732 | 0.083 |
| (a*d) | age.div.plant | −0.046 | −0.123 — 0.030 |  | 0.039 | −1.186 | 0.235 |
| (c*f)+(a*d)+(e*g) | age.div.total | 0.137 | −0.051 — 0.326 |  | 0.096 | 1.428 | 0.153 |
| (c*f*g) | age.obs.env | −0.064 | −0.129 — 0.001 |  | 0.033 | −1.916 | 0.055 |
| (e*g) | age.obs.isolated | 0.235 | 0.033 — 0.437 | * | 0.103 | 2.280 | 0.023 |
| (a*d*g) | age.obs.plant | −0.058 | −0.150 — 0.034 |  | 0.047 | −1.241 | 0.214 |
| (a*d*g)+(c*f*g)+(e*g) | age.obs.total | 0.113 | −0.074 — 0.300 |  | 0.095 | 1.186 | 0.235 |
| age | env | −0.258 | −0.433 — −0.083 | ** | 0.089 | −2.893 | 0.004 |
| (f*g) | env.ind | 0.248 | 0.065 — 0.431 | ** | 0.093 | 2.651 | 0.008 |
| obs.mos | mos.div | 1.247 | 0.691 — 1.803 | *** | 0.284 | 4.397 | <0.001 |
| age | obs.mos | 0.189 | 0.002 — 0.375 | * | 0.095 | 1.983 | 0.047 |
| env | obs.mos | 0.199 | 0.023 — 0.375 | * | 0.09 | 2.213 | 0.027 |
| pc.plant | obs.mos | −0.105 | −0.275 — 0.064 |  | 0.086 | −1.217 | 0.224 |
| age | pc.plant | 0.442 | 0.296 — 0.587 | *** | 0.074 | 5.950 | <0.001 |
| (d*g) | plant.ind | −0.131 | −0.333 — 0.070 |  | 0.103 | −1.276 | 0.202 |
| * p < .05; ** p < .01; *** p < .001 | | | | | | | |

Supplementary table 4: Summary of the path analysis of the effects of time since deglaciation, vascular plant cover and substrate diversity on bryophyte ecological dispersion and diversity (model not used).

| Model Significance | | | |  |  |  |  |
| --- | --- | --- | --- | --- | --- | --- | --- |
| **N** | **χ^2^** | **df** | **p** |  |  |  |  |
| 106 | 0.287 | 3 | 0.963 |  |  |  |  |
| Model Fit | | | | | | |  |
| **CFI** | **RMSEA** | **90% CI** | **TLI** | **SRMR** | **AIC** | **BIC** |  |
| 1 | 0 | 0.000 — 0.000 | 1.055 | 0.01 | 1053.882 | 1093.834 |  |
| Regression Paths | | | | | | | |
| **Predictor** | **DV** | **Standardized** | | | | | |
|  |  | **β** | **95% CI** | **sig** | **SE** | **z** | **p** |
| (c*f) | age.div.env | −0.047 | −0.102 — 0.008 |  | 0.028 | −1.667 | 0.096 |
| (a*d) | age.div.plant | −0.065 | −0.157 — 0.027 |  | 0.047 | −1.381 | 0.167 |
| (c*f)+(a*d)+(e*g) | age.div.total | 0.152 | −0.034 — 0.339 |  | 0.095 | 1.601 | 0.109 |
| (c*f*g) | age.obs.env | −0.063 | −0.128 — 0.002 |  | 0.033 | −1.914 | 0.056 |
| (e*g) | age.obs.isolated | 0.264 | 0.053 — 0.474 | * | 0.107 | 2.458 | 0.014 |
| (a*d*g) | age.obs.plant | −0.088 | −0.201 — 0.026 |  | 0.058 | −1.513 | 0.130 |
| (a*d*g)+(c*f*g)+(e*g) | age.obs.total | 0.113 | −0.072 — 0.298 |  | 0.094 | 1.194 | 0.233 |
| age | env | −0.258 | −0.433 — −0.083 | ** | 0.089 | −2.893 | 0.004 |
| (f*g) | env.ind | 0.245 | 0.064 — 0.427 | ** | 0.093 | 2.644 | 0.008 |
| obs.mos | mos.div | 1.353 | 0.703 — 2.004 | *** | 0.332 | 4.077 | <0.001 |
| age | obs.mos | 0.195 | 0.003 — 0.387 | * | 0.098 | 1.988 | 0.047 |
| env | obs.mos | 0.181 | 0.011 — 0.352 | * | 0.087 | 2.081 | 0.037 |
| plant.cover | obs.mos | −0.122 | −0.293 — 0.048 |  | 0.087 | −1.408 | 0.159 |
| age | plant.cover | 0.529 | 0.401 — 0.656 | *** | 0.065 | 8.141 | <0.001 |
| (d*g) | plant.ind | −0.166 | −0.375 — 0.044 |  | 0.107 | −1.549 | 0.121 |
| * p < .05; ** p < .01; *** p < .001 | | | | | | | |

Supplementary table 5: Summary of the path analysis of the effects of time since deglaciation, vascular plant functional divergence and substrate diversity on bryophyte ecological dispersion and diversity (model not used).

| Model Significance | | | |  |  |  |  |
| --- | --- | --- | --- | --- | --- | --- | --- |
| **N** | **χ^2^** | **df** | **p** |  |  |  |  |
| 106 | 0.742 | 3 | 0.863 |  |  |  |  |
| Model Fit | | | | | | |  |
| **CFI** | **RMSEA** | **90% CI** | **TLI** | **SRMR** | **AIC** | **BIC** |  |
| 1 | 0 | 0.000 — 0.086 | 1.056 | 0.016 | 1086.066 | 1126.018 |  |
| Regression Paths | | | | | | | |
| **Predictor** | **DV** | **Standardized** | | | | | |
|  |  | **β** | **95% CI** | **sig** | **SE** | **z** | **p** |
| (c*f) | age.div.env | −0.046 | −0.103 — 0.011 |  | 0.029 | −1.588 | 0.112 |
| (a*d) | age.div.plant | 0.003 | −0.027 — 0.033 |  | 0.015 | 0.196 | 0.845 |
| (c*f)+(a*d)+(e*g) | age.div.total | 0.128 | −0.056 — 0.311 |  | 0.094 | 1.365 | 0.172 |
| (c*f*g) | age.obs.env | −0.061 | −0.126 — 0.003 |  | 0.033 | −1.867 | 0.062 |
| (e*g) | age.obs.isolated | 0.171 | −0.019 — 0.360 |  | 0.097 | 1.767 | 0.077 |
| (a*d*g) | age.obs.plant | 0.004 | −0.036 — 0.043 |  | 0.02 | 0.196 | 0.844 |
| (a*d*g)+(c*f*g)+(e*g) | age.obs.total | 0.113 | −0.072 — 0.299 |  | 0.095 | 1.196 | 0.232 |
| age | env | −0.258 | −0.433 — −0.083 | ** | 0.089 | −2.893 | 0.004 |
| (f*g) | env.ind | 0.238 | 0.053 — 0.423 | * | 0.094 | 2.525 | 0.012 |
| obs.mos | mos.div | 1.334 | 0.601 — 2.067 | *** | 0.374 | 3.569 | <0.001 |
| age | obs.mos | 0.128 | −0.037 — 0.293 |  | 0.084 | 1.524 | 0.128 |
| env | obs.mos | 0.179 | −0.002 — 0.360 |  | 0.092 | 1.933 | 0.053 |
| plant.fd | obs.mos | 0.014 | −0.126 — 0.154 |  | 0.071 | 0.197 | 0.844 |
| age | plant.fd | 0.211 | 0.031 — 0.391 | * | 0.092 | 2.302 | 0.021 |
| (d*g) | plant.ind | 0.019 | −0.167 — 0.205 |  | 0.095 | 0.197 | 0.844 |
| * p < .05; ** p < .01; *** p < .001 | | | | | | | |
